# E(Q)AGNN-PPIS: Attention Enhanced Equivariant Graph Neural Network for Protein-Protein Interaction Site Prediction

**DOI:** 10.1101/2024.10.06.616807

**Authors:** Animesh, Rishi Suvvada, Plaban Kumar Bhowmick, Pralay Mitra

**Author notes:** (Corresponding Author: Animesh).

## Abstract

Identifying protein binding sites, the specific regions on a protein’s surface where interactions with other molecules occur, is crucial for understanding disease mechanisms and facilitating drug discovery. Although numerous computational techniques have been developed to identify protein binding sites, serving as a valuable screening tool that reduces the time and cost associated with conventional experimental approaches, achieving significant improvements in prediction accuracy remains a formidable challenge. Recent advancements in protein structure prediction, notably through tools like AlphaFold, have made vast numbers of 3-D protein structures available, presenting an opportunity to enhance binding site prediction methods. The availability of detailed 3-D structures has led to the development of Equivariant Graph Neural Networks (GNNs), which can analyze complex spatial relationships in protein structures while maintaining invariance to rotations and translations. However, current equivariant GNN methods still face limitations in fully exploiting the geometric features of protein structures. To address this, we introduce E(Q)AGNN-PPIS ^1^, an Equivariant Attention-Enhanced Graph Neural Network designed for predicting protein binding sites by leveraging 3-D protein structure. Our method augments the Equivariant GNN framework by integrating an attention mechanism. This attention component allows the model to focus on the most relevant structural features for binding site prediction, significantly enhancing its ability to capture complex spatial patterns and interactions within the protein structure. Our experimental findings underscore the enhanced performance of E(Q)AGNN-PPIS compared to current state-of-the-art approaches, exhibiting gains of 8.33% in the Area Under the Precision-Recall Curve (AUPRC) and 10% in the Matthews Correlation Coefficient (MCC) across benchmark datasets. Additionally, our method demonstrates robust generalization across proteins with varying sequence lengths, outperforming baseline methods.

## I. Introduction

**P**ROTEINS, as the primary functional units in biological systems, are integral to the maintenance of structure and function in living organisms [1]. Comprehending their three-dimensional structure is critical for elucidating the underlying mechanisms of diverse biological processes, ranging from enzymatic catalysis, signal transduction and immune response to drug design. A critical challenge in protein structural modeling is the accurate prediction of protein binding sites, also referred to as protein-protein interaction (PPI) site prediction. This problem is of significant importance, as identifying protein binding sites facilitates a deeper understanding of their biological functions [2], subsequently enabling more efficient drug discovery and development processes [3]. Traditionally, protein structural modeling has relied on experimental techniques such as X-ray crystallography, cryo-electron microscopy, and nuclear magnetic resonance for determining protein complex structures [4]. However, these experimental biological methods are often characterized by high resource requirements, substantial time investments, and significant costs. These limitations have motivated the exploration of alternative approaches, particularly computational methods, which have recently demonstrated considerable promise in addressing these challenges.

### Methods for PPI Site Prediction

Accurate binding site prediction necessitates the effective integration of physical, chemical, and geometric information. Recent advancements in structural biology, propelled by the merging of increased processing capacity and innovative learning techniques, have led to the development of various methods for protein-protein interaction prediction [5]–[8]. These learning based methods can be broadly categorized into two groups based on their approach to protein data processing: 1. Machine Learning (ML) and 2. Deep Learning (DL) based methods. ML-based methods typically use protein sequence and structural data, including raw sequences, evolutionary insights, and secondary structure information. These methods employ various ML algorithms, including the Naïve Bayes Classifier [9], Random Forest [5], Support Vector Machines (SVM) [10], and shallow neural networks [11], [12]. Despite the success of ML methods as majority in PPI site prediction, their inability to capture latent structural information often results in sub-optimal performance. The integration of DL methods, such as Convolutional Neural Networks (CNNs) [13], Recurrent Neural Networks (RNNs) [14], and Graph Neural Networks (GNNs) [15], has substantially enhanced the performance over traditional ML methods.

### CNN and RNN Based Methods

CNN-based approaches [16]–[20] have been prominently employed to extract both local and global features from protein sequences and structures. For example, DeepPPIS [16] utilizes TextCNN to distill global features from protein sequences, whereas DeepSite [17] leverages a voxel-based 3-D CNN specifically for binding site prediction. Although CNN-based methods have demonstrated effectiveness in extracting features from proteins, they often overlook long-range dependencies in protein sequences. To overcome this limitation, DLPred [21] introduces an speedup version of the Long Short-Term Memory (LSTM) network, tailored to process sequence-based features derived from the Position-Specific Scoring Matrix (PSSM), alongside physical and physicochemical properties. Furthermore, DELPHI [18] and HN-PPISP [19] employ both Recurrent Neural Networks (RNNs) and CNNs to capture local and long-range features simultaneously. However, conventional convolutional networks face challenges in accurately identifying binding sites, potentially due to the irregularity of protein structures and the arbitrary rotation and translation of proteins in space [22]. Moreover, sequential input to Deep Learning methods is limited by the user-provided features and fails to fully exploit the hidden 3-D structure of proteins during information encoding. In response to these challenges, from the advent of AlphaFold [23], offering accurate complex protein structure predictions, has shifted focus towards methods utilizing tertiary structure information. Consequently, Graph Neural Network (GNN)-based methods have emerged as a robust solution, adept at capturing intricate structural details and enhancing binding site prediction accuracy.

### Graph Neural Network Based Methods

GNN based methodologies are primarily divided into two categories based on their architecture and learning approaches: 1. Traditional GNNs and 2. Geometric GNNs. Traditional GNNs process graph-structured data by progressively refining node representations via a message-passing mechanism among adjacent nodes. For instance, GraphPPIS [6], utilizing a Graph Convolutional Neural Network (GCN) [24], frames the PPI binding site prediction as a node classification task and RGN [25] extends this by incorporating attention mechanism. This approach significantly surpasses earlier deep learning methods by accurately identifying interacting vs. non-interacting amino acid residues. Furthermore, AGAT-PPIS [7] extends the Graph Attention Network (GAT) [26] by incorporating edge features, thus strengthening structural awareness and enhancing translational and rotational invariance. Another noteworthy advancement is GHGPR-PPIS [8], it enhances the traditional GCN framework by integrating an edge-self attention mechanism and employing Heat Kernel techniques to amplify the efficacy of low-frequency filters. To date, GHGPR-PPIS stands as the most effective method for predicting PPI binding sites.

### Limitations of Traditional GNN Methods and the Need for Equivariant Geometric GNNs

Despite the success of traditional GNN methods [6]–[8], [25] on PPI site prediction task, these methods often struggle to capture the intricate geometric dependencies present in protein structures, as they primarily rely on the graph topology and node features while neglecting the crucial spatial or geometric information. Traditional GNNs do not possess built-in equivariance properties in their architecture, making their predictions sensitive to rotations and translations of the input protein structures, which can lead to inconsistent results. Rotation and translation invariance properties are essential for proteins because their function is determined by their 3-D structure, which remains the same regardless of orientation or position in space. For instance, an enzyme’s active site must be recognized irrespective of how the protein is oriented.

To overcome these limitations, equivariant Graph Neural Network (GNN) approaches, such as E(Q)GNN [27] and GVP-GNN [28], have been proposed. In equivariant Graph Neural Networks, 3-D coordinates are crucial as they capture the spatial relationships between atoms or residues. For example, in protein-protein docking, the precise 3-D arrangement of surface residues determines whether two proteins can bind, which cannot be inferred from sequence or graph topology alone [29]. These methodologies are constructed to be invariant under Euclidean group transformations, which encompass rotations, reflections, and translations, in addition to permutations. This characteristic renders them exceptionally apt for the modeling of molecular structures.

In this study, we aim to enhance the identification of binding sites through the application of geometric deep learning techniques. We advocate for the adoption of equivariant GNNs, with a specific focus on augmenting the GVP-GNN architecture [28] through the integration of an attention mechanism. This approach exploits the capabilities of equivariant message passing while handling both scalar and vector features, thus enabling the detailed capture of geometric information present in protein structures. The main contributions of our work are outlined as follows:

- We introduce E(Q)AGNN-PPIS, an equivariant geometric graph neural network architecture that leverages geometric information, designed with a focus on PPI site prediction but potentially applicable to other structural graphs as well. E(Q)AGNN-PPIS is the first method to leverage the expressive power of equivariant message passing, incorporating both scalar and vector features, while introducing an attention mechanism to selectively focus on the most relevant features and interactions during message passing in PPI site prediction task.
- We demonstrate the superior performance of E(Q)AGNN-PPIS on the PPI site prediction task, surpassing state-of-the-art methods. Our model achieves significant improvements in terms of precision (5.45%), recall (9.68%), auprc (8.33%) and F1-score (6.90%), highlighting its effectiveness in accurately identifying binding sites.
- We conduct extensive ablation studies and present visualizations to gain insights into the inner workings of E(Q)AGNN-PPIS. The findings reveal the importance of the attention mechanism and the equivariant message passing in capturing the geometric intricacies of protein structures.

The rest of the manuscript is structured as follows: Section II introduces the fundamental concepts that inspire our method and datasets for training and evaluation. Section III presents E(Q)AGNN-PPIS, our proposed method, and evaluation strategies. Section IV details about experimental setup with results, discussion and ablation studies, followed by conclusion in section V.

## II. Materials and Definitions

In this section, we present a comprehensive overview of the dataset employed in our study, along with a detailed description of the data representation tailored to the problem at hand. To facilitate a clear understanding of the problem and its intricacies, we also provide the necessary definitions and notations.

### A. Datasets Utilized

In order to guarantee an equitable comparison and thorough assessment of the proposed approach, we employ the widely-used benchmark dataset introduced in the state-of-the-art study GHGPR-PPIS [8]. This dataset, which is derived originally from GraphPPIS [6] and refined further, has been extensively utilized in various studies and serves as a standard for evaluating the performance of PPI site prediction methods. The dataset consists of many subsets: a training set labelled as Train_335-1, and three test sets named Test_315-28, Test_60-0, and Ubtest_31-6. Every subset in the dataset is skewed towards negative samples (non-binding sites), making the classification problem challenging. The Train_335-1 and Test_60-0 subsets were principally utilised as benchmark datasets for training and evaluating the proposed model in this work. The other two subsets, Test_315-28 and Ubtest_31-6, were employed to evaluate the model’s ability to generalize to new, unseen data. The details of the datasets used in this study is provided in table I. The extensive details about the datasets can be found in the supplementary dataset section.

**TABLE I.** Descriptive Statistics of Datasets used in this Study derived from GHGPR-PPIS [8].

| Dataset | Protein Complexes | BS<sup>a</sup><br>Count | Non-BS<sup>b</sup><br>Count | Excluded<sup>c</sup><br>Chains | Imbalance<sup>d</sup><br>Ratio |
| --- | --- | --- | --- | --- | --- |
| Train_335-1 | 209 | 10,336 | 58,872 | 1 | 1:6 |
| Test_60-0 | 58 | 2,075 | 11,069 | 0 | 1:5 |
| Test_315-28 | 217 | 8,566 | 51,810 | 28 | 1:6 |
| Ubtest_31-6 | 25 | 711 | 5,206 | 6 | 1:7 |

<sup>a</sup> **BS:** Binding Sites, where amino acids represents the binding sites.
**BS Count:** Total number of true binding sites present in the dataset.
<sup>b</sup> **Non-BS Count:** Total number of non-binding sites present in the dataset.
<sup>c</sup> **Excluded Chains:** Number of removed chains that doesn’t align with their corresponding structure from original dataset [6].
<sup>d</sup> **Imbalance Ratio:** Ratio of binding sites and non-binding sites present in the dataset.

### B. Geometric Graph Construction and Input features

#### 1) Protein Geometric Graph Representation

In this work, we represent each protein structure input, consisting of sequence of *n* amino acids, as undirected graph 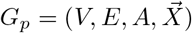.Each graph incorporates both scalar and vector features to describe the 3-D structure of proteins. The node feature matrix *V* has rows 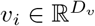 for each of the *N*_*v*_ nodes, where *v*_*i*_ represents the feature vector of node *i*. The edge feature matrix *E* contains vectors 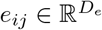 for each edge between nodes *i* and *j*, where an edge exists if *A*_*ij*_ = 1 in the adjacency matrix 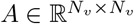 of graph 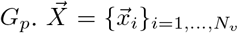 are the positions of central node atoms *C*_*α*_ of amino acids in 3-D Euclidean space, and 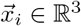.

Each amino acid in the protein sequence is represented as a node in the graph *G*_*p*_. An amino acid is made up of four *backbone atoms*^2^ and a set of side-chain atoms, which are located in 3-D Euclidean space. The construction of each edge *e*_*ij*_ is based on a threshold Euclidean distance between the central carbon atom *C*_*α*_ of the corresponding amino acids represented as node *i* and *j*. The 3-D coordinates of the *C*_*α*_ atom are extracted from the protein’s PDB file. The Euclidean distance *d* is computed as 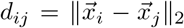 for every pair of *C*_*α*_ atoms in the protein chain’s amino acids. By this we represent a protein chain as an adjacency matrix *A*, where:

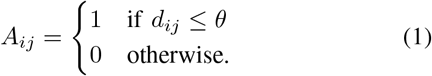

Here *θ* denotes distance threshold, considered as 14Å (angstroms) following the works [6]–[8]. This approach explicitly enforces the structural locality as an inductive bias because nodes farther apart generally does not participate in the interaction. However, stacking multiple GNN layers enables capturing long range interactions as well. We also capture the effect of varying distance threshold as a part of ablation study IV-D.

This study focuses on the utilization of scalar node and edge features which are consistent with the state-of-the-art method GHGPR-PPIS [8]. However we also introduce edge vector features which will be discussed in this section.

#### 2) Node Features

The node features of the geometric protein graph integrate both sequential and structural information of protein chains, offering a comprehensive 62-dimensional feature vector for each amino acid residue. Sequence-based features are derived from Position-Specific Scoring Matrices (PSSM) and Hidden Markov Model (HMM) profiles. PSSM profiles are generated utilizing PSI-BLAST [30], which is used to query the sequence against the UniRef90 database [31], performing three iterations with an E-value of 0.001, while HMM profiles are constructed via HHblits [32], each yielding a feature matrix with dimensions *N*_*v*_ × 20, where *N*_*v*_ represents the length of protein sequence. These matrices are normalized to ensure uniform data scaling across various input sets. Structural features are sourced from the Dictionary of Secondary Structure of Proteins (DSSP) [33], providing a 14-dimensional secondary structural vector per residue. This includes a nine-dimensional^3^ one-hot encoded matrix representing secondary structure categories. An additional four dimensions represent the torsion angles *ϕ* and *ψ*, transformed into sine and cosine values, enhancing rotational invariance. The last dimension quantifies relative solvent accessibility, computed from solvent accessible surface areas. Each amino acid residue is characterised by 7 atomic features, which include atomic mass, B-factor, atom type (side chain or backbone), electrostatic charge, number of bonded hydrogen atoms, presence of aromatic residue, and Van der Waals radius of the atom. These features provide detailed information at the atomic level, which is essential for accurately modelling protein interactions. Finally, the pseudo-position embedding feature (PPEF) encodes the relative positional information of each amino acid residue, benchmarked against a reference residue which we took as the first residue of the protein chain, captures spatial relationships critical for understanding residue interactions.

#### 3) Edge Features

Each edge *e*_*ij*_ in the set of edges, where *i* ≠ *j*, represents a connection between two amino acid residues in the graph and have the following associated features:

- To encode the direction, we calculate the unit vector in the direction from 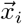 to 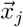 by normalizing the vector 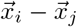.This unit vector serves as a directional feature for the edge connecting center *C*_*α*_ atoms for node *i* and *j*.
- We encode the Euclidean distance between the residues 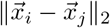 in terms of Gaussian radial basis function.^4^
- The cosine of the angle formed by the vector connecting the residues and a reference vector pointing towards the first amino acid residue of the protein chain.

We summarize all the features used in this study in table II with their corresponding dimensions.

**TABLE II.** Summary of node and edge features used in the E(Q)AGNN-PPIS model and their dimensions.

| Feature Type | Dimension |
| --- | --- |
| PSSM | $N_v \times 20$ |
| HMM | $N_v \times 20$ |
| Atomic Features | 7 |
| DSSP | 14 |
| Pseudo-Position Embedding Feature (PPEF) | 1 |
| Euclidean Distance (Edge) as Gaussian RBFs | 16 |
| Cosine of Angle (Edge) | 1 |
| Unit Vector to encode direction (Edge) | 3 |

**Note:** Node and Edge features are separated by a horizontal line.

### C. Message Passing Neural Networks (MPNNs)

Message Passing Neural Networks (MPNNs) [34] extend the concept of Graph Neural Networks (GNNs) [35], [36] by providing a more general framework for learning representations of graphs. MPNNs define a parameterized function that maps graph structures to feature vectors, operating at both node and graph levels. This approach enables more flexible and expressive modeling of graph-structured data. Since MPNNs [34] employ shared learnable layers across all nodes in the graph, they maintain permutation equivariance, ensuring that the output remains consistent under any reordering of the input nodes.

However, when dealing with geometric graphs, such as those representing protein structures, additional challenges arise. Standard GNNs directly incorporating atomic 3-D coordinates as scalar features would disrupt the equivariance under Euclidean transformations of the input system. Consequently, the primary challenge for Geometric GNNs is to develop an effective method to encode and process geometric graphs while preserving geometric symmetries such as translation invariance, rotational equivariance and reflection equivariance. At a higher abstraction level, the transformation of scalar and vector features from layer *t* to *t* + 1 is facilitated through learnable message and update functions, denoted as MSG and UPD respectively, coupled with a fixed permutation-invariant operator ⊕ (such as mean or sum):

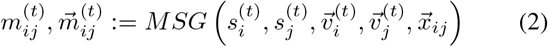

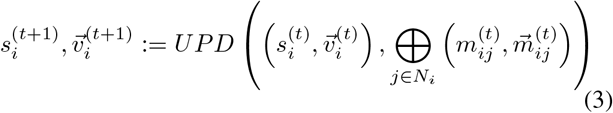

with scalar messages 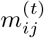 and vector messages 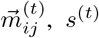 denotes scalar features, 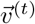 denotes vector features representing geometric features such as 3-D positional vector, and 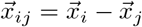 denoting relative position vector.

### D. Invariance and Equivariance

To fully understand the importance of geometric symmetries in MPNNs and Geometric GNNs, it’s crucial to define two key properties: invariance and equivariance. These properties are fundamental to designing models that can effectively handle geometric transformations of input data.

#### Definition II.1 (Invariance)

*Let f* : 𝒢 → 𝒴 *be a function that maps a graph G* ∈ 𝒢 *to an output Y* ∈ 𝒴. *The function f is said to be invariant to a transformation T* : 𝒢 → 𝒢 *if for any graph G* ∈ 𝒢, *f*(*T* (*G*)) = *f*(*G*). *In other words, the output of the function remains unchanged under the transformation T*.

#### Definition II.2 (Equivariance)

*Let f* : 𝒢 → 𝒴 *be a function that maps a graph G* ∈ 𝒢 *to an output Y* ∈ 𝒴, *and let T*_*𝒢*_ : 𝒢 → 𝒢 *and T*_*𝒴*_ : 𝒴 → 𝒴 *be transformations on the input and output spaces, respectively. The function f is said to be equivariant to transformations T*_*𝒢*_ *and T*_*𝒴*_ *if for any graph G* ∈ 𝒢, *f*(*T*_*𝒢*_ (*G*)) = *T*_*𝒴*_ (*f*(*G*)). *In other words, the transformation of the output is equivalent to applying the function to the transformed input*.

In the context of geometric graphs, these properties are particularly important. Let 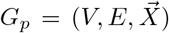 be a geometric graph with 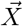 being 3-D geometric vectors (positions, velocities, etc) that are steerable i.e. they transform predictably under the actions of the *E*(3) group (rotations/translations/reflections), and *h* being non-steerable features of nodes *V*. For a geometric graph neural network function *f* to be *E*(3)-equivariant, it must satisfy: 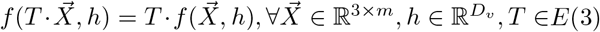. Similarly, *f* is invariant if, 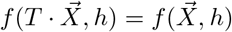.The group action · is instantiated as 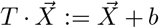 for translation *b* ∈ ℝ^3^ and 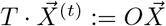. for rotation/reflection *O* ∈ ℝ^3*×*3^.

### E. Problem Statement

Let’s define the binding criterion *C*(*v*_*i*_) for a node *v*_*i*_ to be classified as binding site or not in a geometric protein graph *G*_*p*_ as:

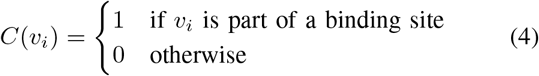

Given this binding criterion and a geometric protein graph *G*_*p*_, our objective is to develop and train an Equivariant Graph Neural Network (GNN) model *f*(*G*_*p*_, *C*) to ensure consistent predictions regardless of the protein’s orientation in 3-D space, which is crucial for accurate binding site identification. This model aims to predict whether any given node *v*_*i*_ in the protein graph *G*_*p*_ is a potential binding site or not. The challenge lies in leveraging the structural and topological information encoded in *G*_*p*_ to effectively classify the nodes, thus enhancing our ability to predict interaction sites in proteins with high accuracy.

## III. Methodology and Evaluation

In this section, we present our proposed method, including it’s mathematical formulation along with evaluation strategies.

### A. E(Q)AGNN-PPIS

#### 1) GVP Embeddings

Geometric Vector Perceptron (GVP-Module) [28] is a simple module to learn scalar and vector valued functions over geometric vectors and scalars. In this work, we employ an enhanced version of the GVP-Module incorporating vector gating [37], which addresses limitations in the original formulation, particularly for atomic-level structure graphs, where individual atoms may not inherently possess orientation information (vector features) at the input level. Vector gating allows the propagation of information from scalar channels into vector channels. The GVP-Module with vector gating [37] as shown in Figure 2, which takes the scalar *s* ∈ ℝ^*n*^ and vector *V* ∈ ℝ^*v×*3^ features as a tuple (*s*; *V*) and returns the new scalar and vector features tuple (*s*^*′*^; *V* ^*′*^) ∈ ℝ^*m*^ × ℝ^*µ×*3^. In Figure 2 *W*_*h*_, *W*_*µ*_, *W*_*m*_ are trainable weight matrices specific to the GVP-Module, *b*_*m*_, *b*_*g*_ are bias vectors, *σ* is a non-linear activation function, and ⊙ denotes element-wise multiplication.

The GVP-Module respects crucial properties for geometric data processing: invariance, equivariance, and expressiveness. It preserves equivariance and invariance under 3-D Euclidean transformations *T*, such that *GV P*(*s, T* (*V*)) = (*s*^*′*^, *T* (*V* ^*′*^)) when *GV P*(*s, V*) = (*s*^*′*^, *V* ^*′*^), as validated in GVP-GNN [28]. The vector gating mechanism enhances the module with universal approximation capability for rotation and reflection equivariant functions *T* : ℝ^*v×*3^ → ℝ^3^ [37].

#### 2) Attention Aggregate Message Passing - AAMP Module

In the proposed E(Q)AGNN-PPIS architecture, the GVP-Module processes both node and edge attributes of the protein graph while preserving equivariance. This processed information is then used in an attention-based mechanism to aggregate data from neighboring nodes. The proposed message passing scheme formulates messages by combining both node and edge features, as detailed in Equation 5. During the aggregation process, the architecture assigns attention scores to these messages from neighboring nodes. These scores effectively weight the contribution of each neighbor, allowing the model to focus on the most relevant information when updating a node’s representation. This approach enables the model to capture complex spatial relationships and interactions within the protein structure, which is essential for accurate prediction of protein-protein interaction sites.

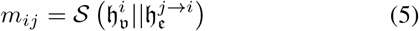

Central to the E(Q)AGNN-PPIS model is the attention mechanism, pivotal for determining the weights of the messages exchanged between nodes in the graph. This mechanism calculates attention scores *α*_*ij*_, which measure the importance of the messages being transmitted from node *j* to node *i*. The adjacency information *A*_*ij*_ is utilized to modulate the attention scores, ensuring that only connected nodes, as defined by the adjacency matrix in Equation 1, contribute to the updated node embeddings. This approach enables the model to focus on the most informative interactions while ignoring irrelevant ones. To calculate the attention scores, we utilize scalar node features, specifically protein node features, as demonstrated in Equations 6 and 7. This design decision enables the model to effectively reflect the intrinsic characteristics of the protein nodes and their impact on the attention mechanism. Once the attention-modulated messages are computed, they are aggregated for each node, effectively combining the weighted contributions from all neighboring nodes.

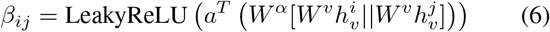

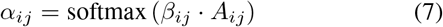

Following the message aggregation step, the node features are updated using Layer Normalization, as shown in Equation 8. This normalization technique is employed to stabilize the learning process by reducing the internal co-variate shift.

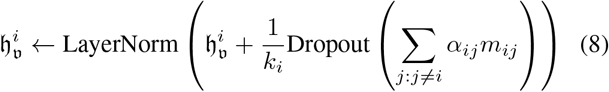

Here, in Equation 5, 𝒮 represents a sequence of three GVP-Module layers. 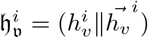 and 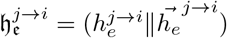 are the embeddings of the node *i* and edge *j* → *i* having both scalar and vector features. *W* ^*α*^, *W* ^*v*^and *a*^*T*^ are learnable parameters metrics of liner layer at different locations and ∥ denotes the concatenation operation. *A*_*ij*_ represents the elements from the adjacency matrix as defined in Equation 1; LeakyReLU serves as the non-linear activation function, characterized by a negative slope of 0.2; *k*_*i*_ denotes the count of incoming messages for node *i*. The LayerNorm [38] is the normalization function applied to each node feature vector independently, ensuring that each feature is normalized based on its own statistics.

Between the propagation steps in the graph, a pointwise feed-forward layer, as shown in equation 9, is employed to update the embeddings of all nodes *i*. This additional layer enables the model to capture more complex interactions between the nodes.

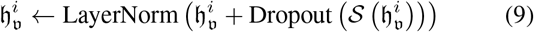

Here 𝒮 at Equation 9 in the sequence of 2 GVP-modules. The graph propagation and pointwise feed-forward layers update not only the scalar but also the vector features for each node in the graph. Our end-to-end architecture, illustrated in Figure 1, employs 8 layers of E(Q)AGNN interconnected via residual connections. This design mitigates the oversmoothing issue while enhancing the capture of structural information.

**Fig. 1.**
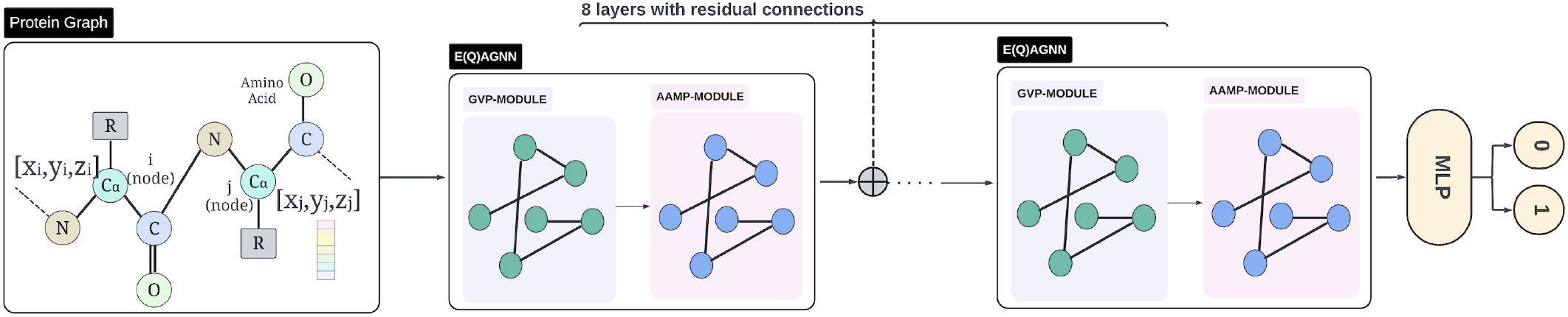
End-to-end architecture of E(Q)AGNN-PPIS for protein-protein interaction site prediction. The model processes a protein graph representation depicting amino acids as nodes with their constituent atoms and 3-D coordinates, through 8 E(Q)AGNN layers, each containing Geometric Vector Perceptron (GVP) and Attention Aggregated Message Passing (AAMP) modules. These layers are connected via residual connections. The final node embeddings are fed into a Multi-Layer Perceptron (MLP) for binary classification of each amino acid as a binding site (1) or non-binding site (0).

**Fig. 2.**
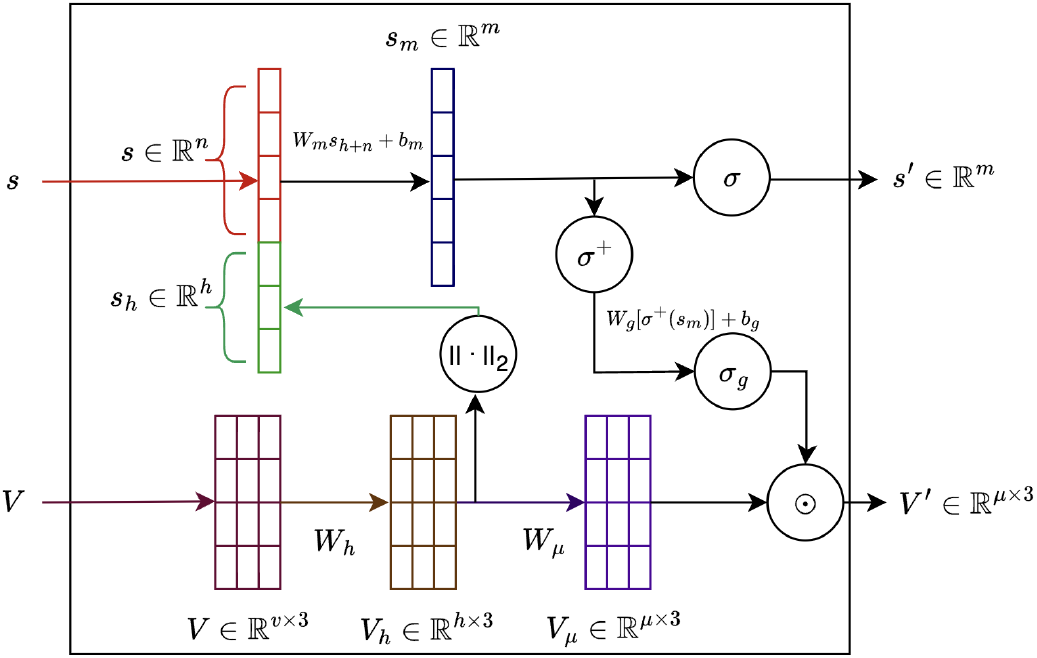
Schematic representation of GVP-Module [37].

#### 3) Building Equivariant Features

In protein structure analysis, we leverage rich atomic-level information to construct equivariant features, distinguishing our approach from typical point cloud datasets. We compute both node and edge vector features, which satisfy specific equivariance conditions (for proof, refer to Appendix C-A and C-B), ensuring our model maintains geometric consistency under spatial transformations. For nodes, we compute vector features 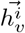 as described in Proposition 1. For edges, we introduce vector features 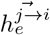 that capture directional information, as detailed in Proposition 2. These vector features are complemented by scalar features for both nodes and edges, derived from various physicochemical properties and sequence-based information (see Sections II-B2 and II-B3). This hybrid approach allows our E(Q)AGNN-PPIS model to simultaneously process both structural and biochemical aspects of protein interactions while maintaining appropriate equivariance properties.

##### Proposition 1.

*Let* 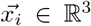 *be the position vector of the i*^*th*^ *node representing an amino acid, and* 𝒩_*i*_ *be the set of neighbors of node i. The node vector feature* 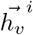 *defined as:*

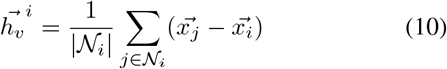

*is invariant under translations and equivariant under the orthogonal group O(3), which includes rotations and reflections*.

##### Proposition 2.

*Let* 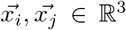 *be the position vectors of nodes i and j. The edge vector feature* 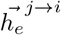 *defined as:*

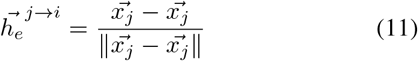

*is invariant under translations and equivariant under the orthogonal group O(3), which includes rotations and reflections*.

#### 4) Model Equivariance

Having introduced all components of our E(Q)AGNN-PPIS model, including the GVP Embeddings, Attention Aggregate Message Passing (AAMP) Module, and the construction of equivariant features, we now establish the equivariance properties of the entire model. The The proposition 3 shows that E(Q)AGNN-PPIS preserves the equivariance properties of its input features throughout its operations.

##### Proposition 3.

*The E(Q)AGNN-PPIS model, including its attention mechanism defined in Equations* (5)*–*(9), *is equivariant under the actions of the orthogonal group O*(3).

*Proof*. See Appendix C-C □

### B. Performance Evaluation Metrics

To evaluate our E(Q)AGNN-PPIS method for predicting protein-protein interaction (PPI) binding sites the metrics like Accuracy, Precision, Recall, F1 Score, Matthews Correlation Coefficient (MCC), and Area Under the Precision-Recall Curve (AUPRC) have been used in this study. This comprehensive approach allows for direct comparison with state-of-the-art methods in PPI site prediction. As evident from Table I, the datasets used in this study exhibit a significant class imbalance, which is inherent to the PPI binding site prediction problem. Consequently, the most informative and commonly used metrics for evaluating model performance in such scenarios are F1 Score, MCC, and AUPRC, as they provide a more comprehensive assessment of the model’s ability to handle class imbalance effectively. The equations to compute these metrics are given in Appendix B.

## IV. Experiments and Results

### A. Implementation Details and Hyper-Parameters

For the development of the E(Q)AGNN-PPIS, we employed the PyTorch v2.2.1[39] and PyTorch-Geometric v2.4.0 [40] libraries. Optimization was performed using the Adam [41] optimizer over 100 epochs with an initial learning rate of 0.0005, targeting the minimization of the Binary Cross-Entropy loss as specified in equation 12. The architecture included eight layers of E(Q)AGNN, incorporating residual connections to counter over-smoothing effects [42]. To reduce overfitting, we applied a dropout rate of 0.1 throughout the training process. A plateau learning rate scheduler was also utilized, adjusting the learning rate based on stagnation in the validation loss. To compensate for class imbalances in the dataset, we adjusted the class weights in the Binary Cross-Entropy loss function to a 1:4 ratio, with the positive class (interaction sites) weighted more heavily than the negative class (non-interaction sites).

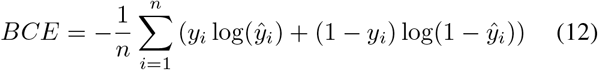

where *n* is the number of samples, and *y*_*i*_ and *ŷ*_*i*_ represent the actual and predicted labels, respectively.

A detailed hyper-parameters search and finally used hyper-parameters for training are detailed in table III.

**TABLE III.**
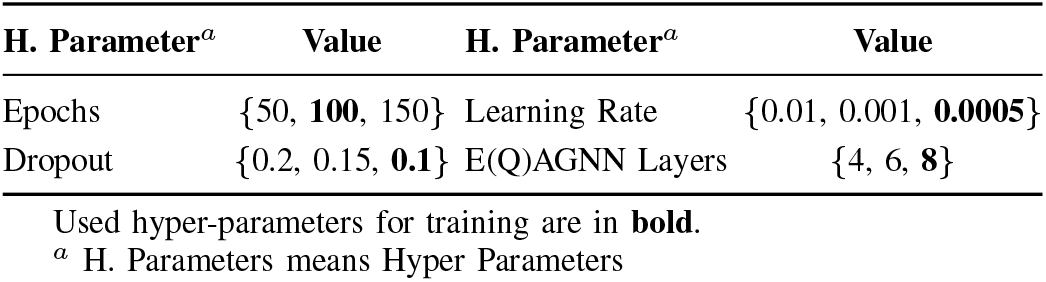
Hyper-Parameter search for E(Q)AGNN-PPIS.

| H. Parameter<sup>a</sup> | Value | H. Parameter<sup>a</sup> | Value |
| --- | --- | --- | --- |
| Epochs | {50, **100**, 150} | Learning Rate | {0.01, 0.001, **0.0005**} |
| Dropout | {0.2, 0.15, **0.1**} | E(Q)AGNN Layers | {4, 6, **8**} |

Used hyper-parameters for training are in **bold**.
<sup>a</sup> H. Parameters means Hyper Parameters

### B. Performance Comparison with Baseline Methods

Our proposed method E(Q)AGNN-PPIS, have been compared with various baseline methods including GHGPR-PPIS [8], which is the state-of-the-art method for the dataset used. In order to make the comparison fare, we conducted a comparative analysis against a spectrum of established baseline methods including machine learning based methods such as PSIVER [9], ProNA2020 [43] which primarily leverages sequential features, and deep learning based methods like DeepPPISP [16], SPPIDER [44], MaSIF-site [45], which use local and global features of protein sequence either by extracting protein molecular surface fingerprints [45] or directly applying CNNs at the residue level [16]. Furthermore, our evaluation included comparisons with established GNN-based methods such as GraphPPIS [6], AGAT-PPIS [7] and GHGPR-PPIS [8]. These methods construct protein graphs from it’s tertiary structures and employ various GNN methods to learn node features: GraphPPIS used GCN [24] with residual connections to capture the local structural information, AGAT-PPIS used GAT [26] with edge features to learn the importance of neighboring nodes and GHGPR-PPIS utilizes GraphHeat [46] with self edge attention mechanism to model the heat diffusion process on the protein graphs. All these methods aim to accurately predict PPI binding sites by exploiting the graph representation of proteins and learning informative node features.

Experimental results presented in Table IV demonstrate the superior performance of our proposed method, E(Q)AGNN-PPIS, in predicting PPI binding sites compared to existing state-of-the-art techniques. E(Q)AGNN-PPIS achieves the highest scores across all evaluation metrics on the test 60 dataset, with an significant improvement of **10%** in MCC (0.55), **6.90%** in F1 score (0.62), **9.68%** in Recall (0.68), **5.45%** in Precision (0.58) and **1.28%** in Accuracy (0.87) when compared to the second best method GHGPR-PPIS. Given the imbalanced nature of the dataset, AUPRC becomes the most crucial metric for evaluation [47]. In this regard, E(Q)AGNN-PPIS demonstrates a significant improvement, achieving a **8.33%** higher AUPRC (0.65) compared to GHGPR-PPIS. The superior performance of E(Q)AGNN-PPIS stems from its implementation of equivariant geometric deep learning with attention mechanism. By incorporating GVP-Module, our architecture preserves crucial 3-D structural information of proteins while processing node and edge features. The attention-based message passing scheme then allows the model to dynamically focus on the most relevant spatial relationships between residues. This combination enables E(Q)AGNN-PPIS to capture both local geometric patterns and long-range interactions critical for accurate PPI site prediction.

**TABLE IV.**
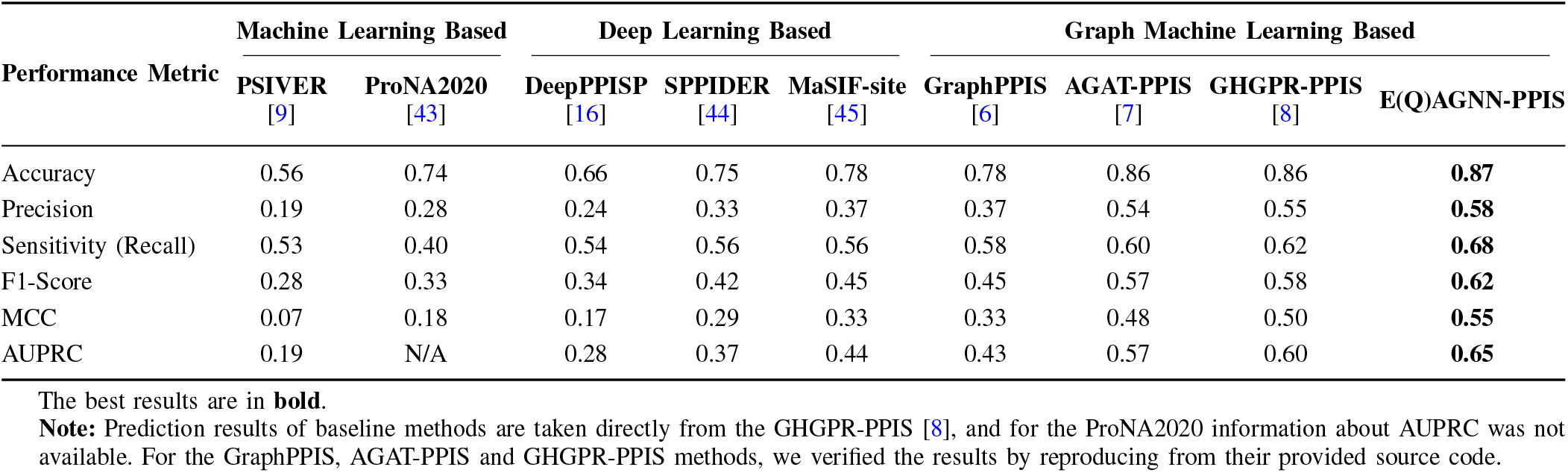
Comparative analysis of E(Q)AGNN-PPIS and other baseline methods on the Test 60 dataset.

### C. Generalization and Real-World Deployment Opportunities for E(Q)AGNN-PPIS in Protein Interaction Studies

#### 1) Generalization

To assess the generalization capability of our proposed method, we conducted evaluations on two independent and diverse test sets, namely Test_315-28 and Ubtest_31-6 [7]. Table V presents a comprehensive comparison of our results with current state-of-the-art (SOTA) methods, including GraphPPIS [6], AGAT-PPIS [7], GHGPR-PPIS [8], and several others. Considering the imbalanced nature of the datasets, AUPRC serves as the crucial evaluation metric [47]. The empirical findings demonstrate the robust generalization ability of our method, as it consistently outperforms both AGAT-PPIS and GHGPR-PPIS on the Ubtest_31-6 dataset across all metrics, notably achieving an **15.34%** improvement in AUPRC over AGAT-PPIS and a **14.71%** improvement over GHGPR-PPIS. Moreover, on the Test_315-28 dataset, our method achieves the highest AUPRC, representing a **5.1%** improvement over AGAT-PPIS and a **6.93%** improvement over GHGPR-PPIS, alongside notable enhancements in both F1 and MCC scores. Furthermore, our method demonstrates superior performance compared to earlier methods such as DeepPPISP, SPPIDER, and MaSIF-site, showcasing significant advancements in the field. These outcomes highlight the effectiveness of our approach in handling unseen data inputs and its potential for real-world applications in protein-protein interaction site prediction.

**TABLE V.** Generalization performance comparison of E(Q)AGNN-PPIS with current SOTA methods.

| Method | Test_315-28 |  |  | Utest_31-6 |  |  |
| --- | --- | --- | --- | --- | --- | --- |
|  | F-1 | MCC | AUPRC | F-1 | MCC | AUPRC |
| DeepPPISP | N/A | 0.169 | 0.256 | N/A | 0.162 | 0.217 |
| SPPIDER | N/A | 0.294 | 0.376 | N/A | 0.222 | 0.260 |
| MaSIF-site | N/A | 0.304 | 0.372 | N/A | 0.141 | 0.225 |
| GraphPPIS | 0.420 | 0.336 | 0.423 | 0.386 | 0.280 | 0.323 |
| AGAT-PPIS | 0.559 | 0.481 | 0.572 | 0.414 | 0.327 | 0.365 |
| GHGPR-PPIS | 0.564 | 0.486 | 0.562 | 0.441 | 0.356 | 0.367 |
| **E(Q)AGNN-PPIS** | **0.573** | **0.497** | **0.604** | **0.444** | **0.362** | **0.421** |

The best results are in **bold**.
**Note:** The generalization outcomes for the methods GraphPPIS, AGAT-PPIS, and GHGPR-PPIS have been reproduced using their respective source codes. The results for DeepPPISP, SPPIDER, and MaSIF-site are referenced from GraphPPIS [6], with the F-1 score data not being available.

#### 2) Real-World Deployment Opportunities for E(Q)AGNN-PPIS

To assess the practical utility of the E(Q)AGNN-PPIS model, we undertook a case study analyzing individual proteins selected from the Test_60 dataset. Specifically, we examined 10 proteins: 3cf4(A), 4cdg(A), 3zhe(B), 1wpx(A), 3fmo(A), 2fhz(B), 3mcb(A), 3vz9(D), 4nqw(B) and 4je3(B), with the letters in parentheses indicating the specific protein chains. We have selected these proteins carefully by choosing 5 longest and 5 shortest proteins from the dataset to represent diversity. We present our observations particularly on the 1wpx(A) protein chain, which is one of the longest in the dataset, consisting of 421 amino acid residues including 46 labelled as binding sites. Detailed observations for the other proteins are documented in Appendix A.

For the protein chain 1wpx(A), we compared the E(Q)AGNN-PPIS with two SOTA models, GHGPR-PPIS and AGAT-PPIS, focusing on their ability to accurately identify binding sites. As per the results summarized in Table VI, the E(Q)AGNN-PPIS identified 33 residues as binding sites, of which 32 were confirmed as true positives and only one as a false positive, with 14 residues as false negative. In contrast, AGAT-PPIS and GHGPR-PPIS predicted significantly less and more binding sites, respectively and both the methods achieved F1-Score of **49%** and **62%**. Notably, our method achieved a superior F1-Score of **81%**, demonstrating its effectiveness in identifying potential binding sites within complex protein chains. The results exhibits the enhanced capability of E(Q)AGNN-PPIS to not only predict with high precision but also to maintain a lower rate of false positives, thus providing a robust tool for the prediction of PPI binding sites. In Figure 3, we present the comparative visualizations of the predictions from the GHGPR-PPIS 3c and E(Q)AGNN-PPIS 3b methods for the protein ‘1wpx(A)’. The results indicate that the predictions made using E(Q)AGNN-PPIS align more closely with the actual observations, showcasing its enhanced accuracy in PPI binding sites prediction.

**TABLE VI.**
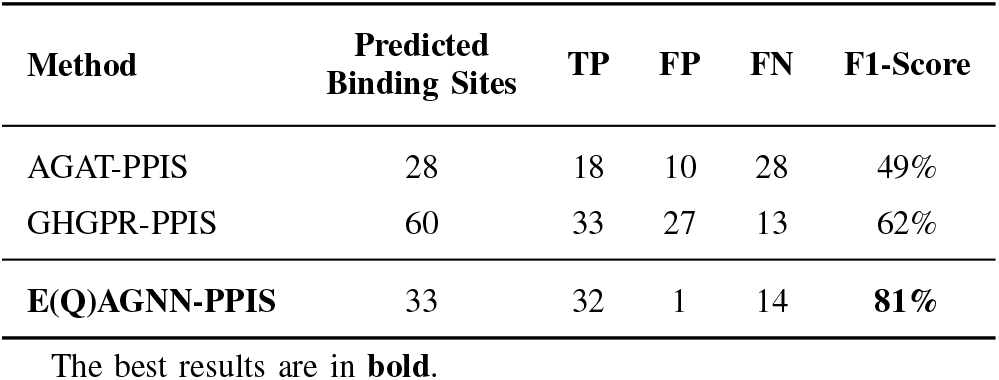
Comparison of predicted results from E(Q)AGNN-PPIS on protein ‘1wpx(a)’ of length 421 with state-of-the-art methods.

| Method | Predicted Binding Sites | TP | FP | FN | F1-Score |
| --- | --- | --- | --- | --- | --- |
| AGAT-PPIS | 28 | 18 | 10 | 28 | 49% |
| GHGPR-PPIS | 60 | 33 | 27 | 13 | 62% |
| **E(Q)AGNN-PPIS** | 33 | 32 | 1 | 14 | **81%** |

The best results are in **bold**.

**Fig. 3.**
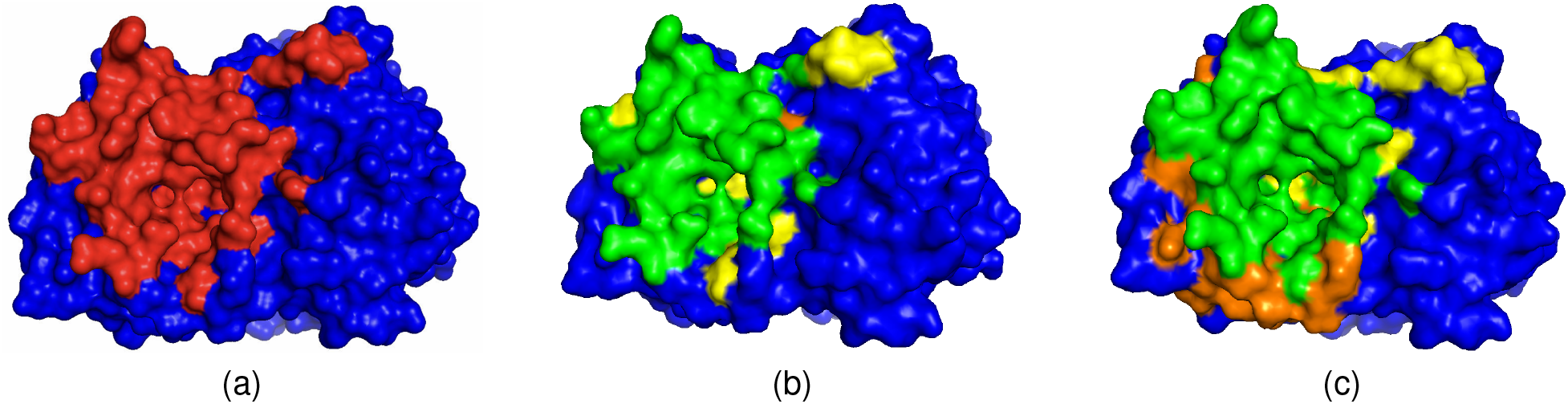
An example case of protein ‘1wpx(A)’, (a) original binding site, (b) prediction using our method E(Q)AGNN-PPIS, and (c) prediction using GHGPR-PPIS. We used color codes Red, Green, Orange, Yellow and Blue to indicate original binding sites, True Positive, False Positive, False Negative and non binding sites, respectively. Visualization is done using PyMOL [48] tool.

#### 3) Length-invariant Performance of E(Q)AGNN-PPIS

To evaluate the effectiveness of E(Q)AGNN-PPIS across varying protein lengths, we categorized the Test_60 dataset into small (52-127 residues), medium (128-247 residues), and large (253-766 residues) proteins. As shown in Table VII, E(Q)AGNN-PPIS consistently outperforms AGAT-PPIS and GHGPR-PPIS across all length categories, achieving average F1 scores of 0.672, 0.634, and 0.563 for small, medium, and large proteins, respectively. This robust performance, with improvements of up to **32.15%** over AGAT-PPIS and **10.3%** over GHGPR-PPIS, demonstrates E(Q)AGNN-PPIS’s length-invariant predictive capability. The model’s ability to maintain high accuracy across diverse protein sizes can be attributed to its effective utilization of geometric equivariance properties and attention mechanisms, making it well-suited for real-world binding site prediction tasks.

**TABLE VII.** Average F1 Scores for Different Protein Lengths Across Models.

| Protein Length | E(Q)AGNN-PPIS | AGAT-PPIS | GHGPR-PPIS |
| --- | --- | --- | --- |
| Small | **0.672** | 0.599 | 0.639 |
| Medium | **0.634** | 0.599 | 0.599 |
| Large | **0.563** | 0.426 | 0.510 |

### D. Ablation Studies

To demonstrate the contribution of each component of our pipeline to the overall performance of PPI binding site prediction, we have performed an ablation study through additional experiments.

#### 1) Evaluating the Effect of Threshold (θ) on protein distance map

As mentioned in section II-B1 we derived the adjacency matrix by using a threshold *θ* on the protein distance matrix; as shown in Equation 1. Here we report the performance of E(Q)AGNN-PPIS at various distance thresholds on Test_60 dataset.

From the Figure 4, it can be seen that as the distance cutoff threshold increases, the model’s performance generally improves across all metrics because information flow increases, until reaching a peak at around 14Å. Beyond this point, the performance metrics experience a slight decline. As the cutoff increases, the protein graph becomes increasingly dense, leading to oversquashing [49], which occurs when long-range dependencies in dense graphs cannot be effectively captured, resulting in information loss. The optimal 14Å cutoff thus represents a balance between sufficient information propagation and preservation of meaningful geometric relationships in the protein structure. The Accuracy and AUPRC exhibit the highest values among the metrics, followed by F1 Score, MCC, Precision, and Recall. Throughout this work we have utilized the distance cutoff of 14Å.

**Fig. 4.**
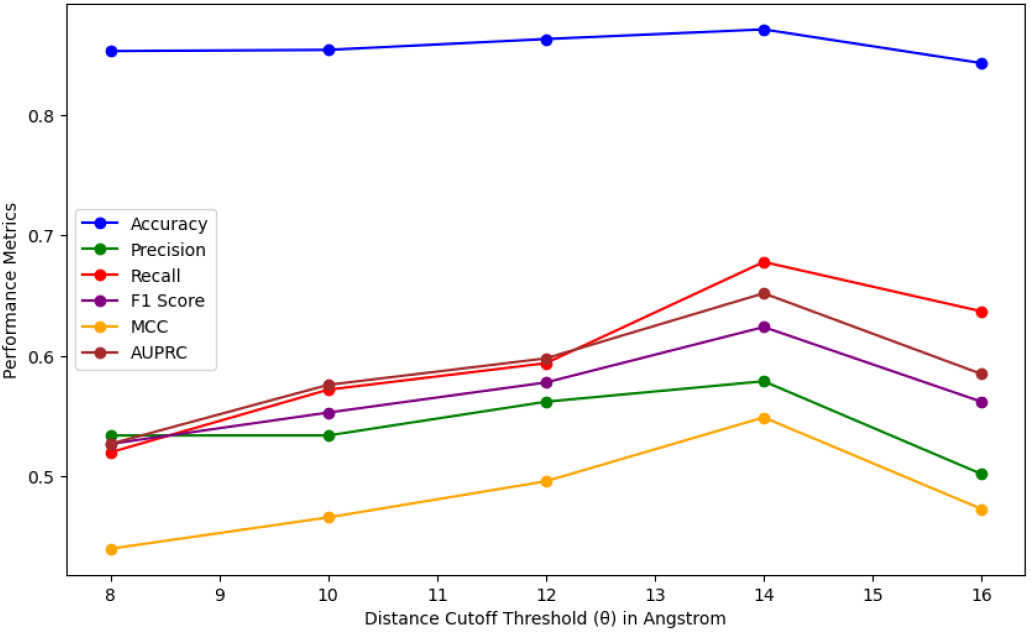
Performance of E(Q)AGNN-PPIS on different distance cutoff threshold *θ*. Various matrix are presented in different colours.

#### 2) Effect of number of Layers

This section investigates the influence of model depth on the performance of the E(Q)AGNN-PPIS model. Figure 5 illustrates the effect of varying the number of E(Q)AGNN layers on a range of evaluation metrics, using the Test_60 dataset. The results demonstrate that increasing the number of layers from 5 to 10 generally enhances the model’s performance, with Accuracy and AUPRC exhibiting the highest values and the most consistent improvement. However, the optimal number of layers is dependent on the specific metric and the trade-off between performance gain and computational complexity. In this study, we employed 8 layers across all experiments, as it provides a favorable balance between performance and computational efficiency.

**Fig. 5.**
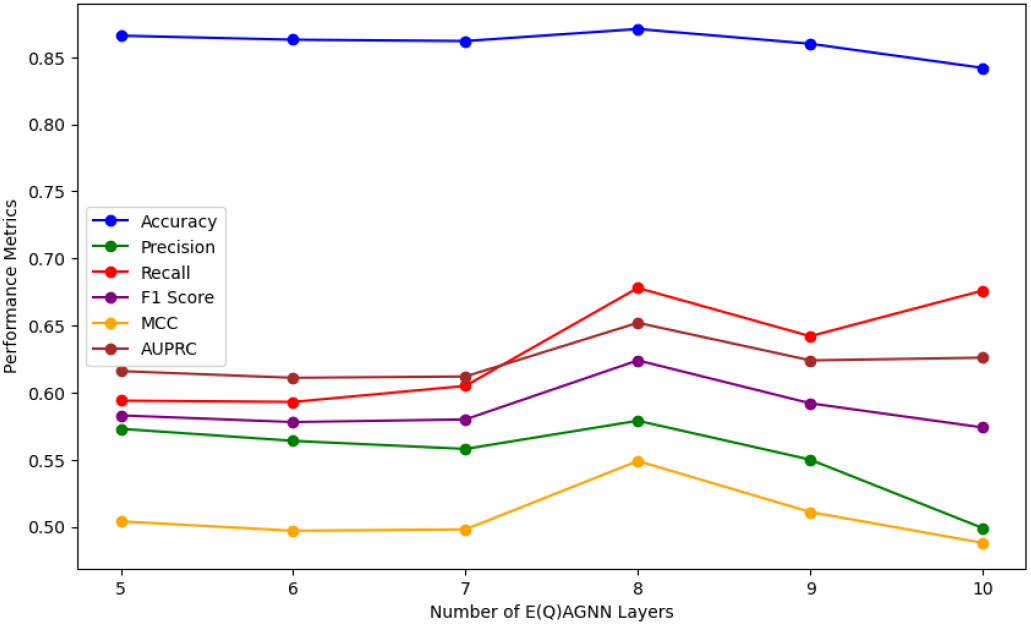
Performance of E(Q)AGNN-PPIS on different number of E(Q)AGNN layers.

## V. Conclusion

In this research, we introduce E(Q)AGNN-PPIS, an attention-enhanced equivariant graph neural network specifically designed for predicting protein binding sites without prior knowledge of binding partners. Our methodology integrates an innovative attention mechanism that processes both scalar and vector features of the protein graph, facilitating effective node representation learning. By incorporating eight layers of E(Q)AGNN with residual connections, our model demonstrates superior robustness and generalization capabilities compared to other state-of-the-art methods across diverse test datasets. While our approach incorporates structural features, evolutionary information, secondary structure level information, and atomic level features, we acknowledge that integrating additional physico-chemical properties (such as dissociation constants and optical activity etc.) could potentially enhance prediction accuracy. Moreover, our current method focuses on unilateral binding site prediction. A natural extension of this work would be to leverage binding partner information to identify complementary binding sites, considering both geometric and physico-chemical compatibility. This approach could pave the way for modeling protein dimeric complex structures, significantly advancing our understanding of protein-protein interactions.

Looking ahead, we plan to expand this work by developing a model that more effectively addresses these limitations while maintaining adherence to the geometric intricacies of protein data. We are optimistic that our approach could be readily adapted to a broader range of applications, including the prediction of protein interactions with small ligands, DNA, and RNA, thereby contributing to the development of a more generalized model for biomolecular interaction prediction.

## Appendix A Ablation Results

Here we provide the full prediction details of selected protein from our method, E(Q)AGNN-PPIS as explained in Section IV-C2 and provide the comparison with the state-of-the-art method GHGPR-PPIS.

**TABLE VIII.** Predicted results from E(Q)AGNN-PPIS and GHGPR-PPIS method on various proteins of different lengths.

| Protein-ID<sup>a</sup> | Length | Method | Predicted binding sites | TP | FP | FN | F1-Score |
| --- | --- | --- | --- | --- | --- | --- | --- |
| 3cf4(A) | 766 | Ours | 112 | 36 | 76 | 21 | 43% |
|  |  | GHGPR-PPIS | 98 | 35 | 63 | 22 | **45%** |
| 4cdg(A) | 625 | Ours | 69 | 21 | 48 | 18 | **39%** |
|  |  | GHGPR-PPIS | 99 | 18 | 81 | 21 | 26% |
| 3fmo(A) | 421 | Ours | 34 | 23 | 11 | 11 | **68%** |
|  |  | GHGPR-PPIS | 25 | 17 | 8 | 17 | 58% |
| 3zhe(B) | 390 | Ours | 42 | 26 | 16 | 11 | **66%** |
|  |  | GHGPR-PPIS | 55 | 26 | 29 | 11 | 57% |
| 3mcb(A) | 54 | Ours | 40 | 23 | 17 | 6 | 67% |
|  |  | GHGPR-PPIS | 48 | 27 | 21 | 2 | **70%** |
| 3vz9(D) | 60 | Ours | 31 | 25 | 6 | 8 | **78%** |
|  |  | GHGPR-PPIS | 33 | 25 | 8 | 8 | 76% |
| 4nqw(B) | 77 | Ours | 39 | 38 | 1 | 6 | **92%** |
|  |  | GHGPR-PPIS | 36 | 32 | 4 | 12 | 80% |
| 4je3(B) | 73 | Ours | 34 | 30 | 4 | 7 | **85%** |
|  |  | GHGPR-PPIS | 44 | 29 | 15 | 8 | 72% |
| 2fhz(B) | 93 | Ours | 43 | 31 | 12 | 1 | **83%** |
|  |  | GHGPR-PPIS | 29 | 16 | 13 | 16 | 52% |

<sup>a</sup> In Protein-ID column, it follows the structure PDB-ID(chain). 'Ours' refers to E(Q)AGNN-PPIS method. Best method values are in **bold**.

## Appendix B Performance Metrics

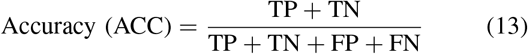

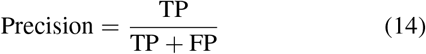

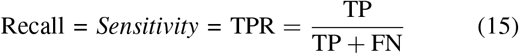

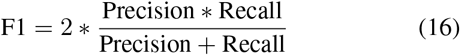

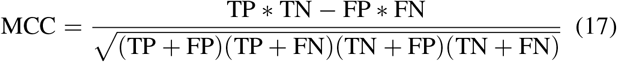

In the above metrics, TP (True Positive) denotes the number of correctly identified PPI binding sites, whereas TN (True Negative) refers to the correct predictions of non-PPI binding sites. Conversely, FP (False Positive) indicates instances where non-binding sites are erroneously predicted as binding sites, and FN (False Negative) marks actual binding sites that are misclassified as non-binding. Additionally, TPR (True Positive Rate), also known as sensitivity, quantifies the ratio of actual PPI binding sites that are accurately detected by the model.

## Appendix C Proofs

### A. Proof of Proposition 1

#### Proposition 1

Let 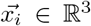 be the position vector of the *i*^*th*^ node representing an amino acid, and 𝒩_*i*_ be the set of neighbors of node *i*. The node vector feature 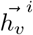 defined as:

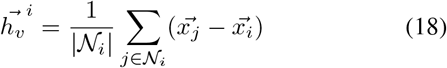

is invariant under translations and equivariant under the orthogonal group O(3).

*Proof*. We prove the invariance under translations and equivariance under O(3) separately:

**Case 1. Translation:** Under a translation 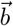,the positions transform as 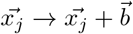 and 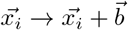. The transformed 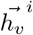 is:

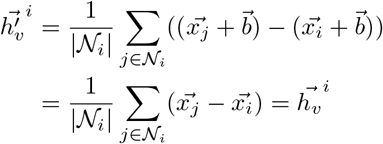

This demonstrates translation invariance of 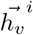.

**Case 2. O(3) Transformation:** For an orthogonal matrix *R* ∈ ℝ^3*×*3^, vectors transform as 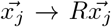 and 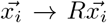 The transformed 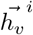 is:

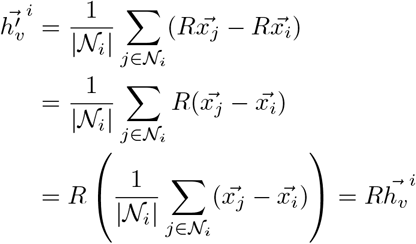

This demonstrates equivariance of 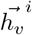 under O(3) transformations. □

### B. Proof of Proposition 2

#### Proposition 2

Let 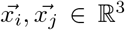 be the position vectors of nodes *i* and *j*. The edge vector feature 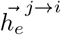 defined as:

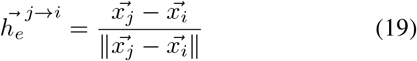

is invariant under translations and equivariant under orthogonal group O(3).

*Proof*. We prove the invariance under translations and equivariance under O(3) separately:

**Case 1. Translation:** Under a translation 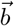,the positions transform as 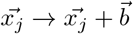 and 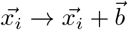. The transformed 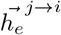 is:

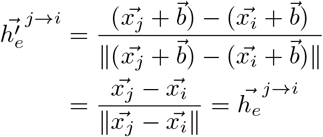

This demonstrates translation invariance of 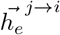.

**Case 2. O(3) Transformation:** For an orthogonal matrix *R* ∈ ℝ^3*×*3^, vectors transform as 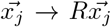 and 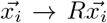. The transformed 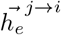is:

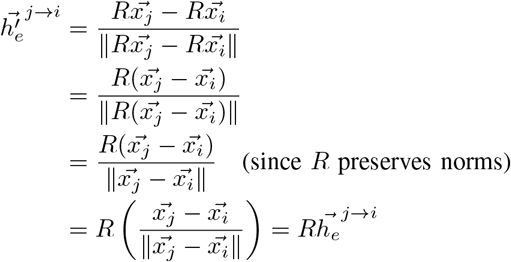

This demonstrates equivariance of 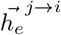 under O(3) transformations. □

### C. Proof of Proposition 3

#### Proposition 3

We proof that the E(Q)AGNN-PPIS model, including its attention mechanism as defined in Equations (5)– (9), is equivariant under the actions of the orthogonal group *O*(3).

*Proof*. Let *R* ∈ ℝ^3*×*3^ be an orthogonal metrix.

**Base case: Input features** The input vector features 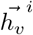 for any node *i* and 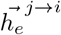 for the edge between nodes *j* and *i* are equivariant under any transformation *R* ∈ *O*(3) as established in Proposition 1, 2 and scalar features are invariant by definition.

**Inductive step:** These input features are used in subsequent layers of our method, we prove that operations in layer *t* + 1 preserve equivariance as input features at layer *t* are equivariant.

1. *GVP-Module:* The GVP-Module preserves equivariance as shown in [37]. For completeness:

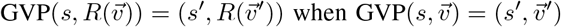
2. *Attention Mechanism:* The attention scores *α*_*ij*_ are computed using only scalar features:

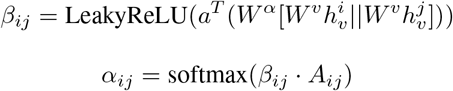 Since scalar features and the adjacency matrix *A*_*ij*_ are invariant under any transformation *R, α*_*ij*_ is also invariant.
3. *Message Passing:* The message function is:

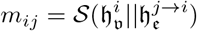

where 𝒮 is a sequence of GVP-Modules. Since GVP-Modules preserve equivariance, and concatenation of equivariant features remains equivariant, *m*_*ij*_ is equivariant.
4. *Feature Update:* The feature update is:

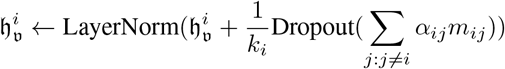

Since *α*_*ij*_ is invariant (scalar), *m*_*ij*_ is equivariant and sum of equivariant features is equivariant, therefore, the updated features 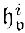 are equivariant.

By induction, all layers of the network preserve equivariance. Therefore, the entire E(Q)AGNN-PPIS model is equivariant under *O*(3) transformations. □

**Animesh** received the B.Tech degree in Electrical and Electronics Engineering from the Pranveer Singh Institute of Technology, Kanpur, India, in 2016 and M.tech degree in Electrical Engineering with specialization in Control System from the National Institute of Technology (NIT), Kurukshetra, India in 2020. He is currently pursuing a Ph.D. in the Department of Artificial Intelligence at the Indian Institute of Technology (IIT), Kharagpur. His research interests include graph machine learning and its applications.

**Rishi Suvvada** pursued an interdisciplinary Dual degree with B.Tech in Chemical Engineering and M.Tech in Artificial intelligence and Machine learning from the Department of Artificial Intelligence at Indian Institute of Technology Kharagpur,India from 2019 to 2024. He is currently pursuing his corporate stint working as a Data Scientist in a Heath Tech startup called Hilabs His research interests include graph machine learning and its applications, AI in medicine and finance space.

**Plaban Kumar Bhowmick** received the B.Tech degree from University of Calcutta, India, 2002, and the master’s and Ph.D. degrees in Computer Science and Engineering from the Indian Institute of Technology (IIT) Kharagpur, India, in 2006 and 2011, respectively. He is an Associate Professor with the Department of Artificial Intelligence, IIT Kharagpur. He has been Co-Principal Investigator and Technical Lead of the National Digital Library of India project. His research areas include text processing, artificial intelligence in education, semantic web technology, knowledge graphs, graph machine learning, and digital library technologies.

**Pralay Mitra** received the BSc degree with honors in physics and BTech degree in Computer Science and Engineering from the University of Calcutta, India, in 1999 and in 2002, respectively. He finished the ME degree in computer science and engineering from the Bengal Engineering and Science University, Shibpur (currently known as IIEST, Shibpur), India, in 2004, and the PhD degree from the Indian Institute of Science, Bangalore, in 2010. He was the research associate with the Indian Institute of Science, Bangalore (from 2010 to 2011) and the research fellow with the University of Michigan, Ann Arbor (from 2011-2013). Since 2013, he has been at the Department of Computer Science and Engineering, Indian Institute of Technology Kharagpur, India as an Assistant Professor (2013-2020) and then as an Associate Professor (2020-present). He is actively working on bioinformatics and computational biology. Over the years, he has developed expertise in modeling and designing protein structures and protein functions. He has developed a number of algorithms for protein-protein docking, predicting protein assembly from the crystal structures, and protein design.

## Footnotes

1 We will release the code upon the decision from reviewers.

2 The atoms *C*_*α*_, *C, N* and *O*. The alpha carbon (*C*_*α*_) is the central carbon atom in every amino acid residue.

3 *H* and *G* represent the alpha-helix and 3_10_ helix, respectively. *I*: -helix and *E* represents an extended beta-strand. *B*: beta-strand, *T* : turns, *S*: bend, The variable *C* represents secondary structures of proteins that are not classified into any specific category, as well as those that are unknown.

4 We employ 16 Gaussian radial basis functions (RBFs) with centers uniformly distributed between 0 and 20 angstroms to effectively capture spatial relationships.

